# Ectodysplasin signaling via Xedar is required for mammary gland morphogenesis

**DOI:** 10.1101/2022.09.08.507158

**Authors:** Abigail R. Wark, Reiko Tomizawa, Daniel Aldea, Blerina Kokalari, Bailey Warder, Yana G. Kamberov

## Abstract

The Ectodysplasin A2 receptor (XEDAR), is a member of the tumor necrosis factor receptor subfamily and is a mediator of the Ectodysplasin (EDA) signaling pathway. EDA signaling plays evolutionarily conserved roles in the development of the ectodermal appendage organ class that includes hair, eccrine sweat glands, and mammary glands. Loss of function mutations in *Eda*, which encodes the two major ligand isoforms, EDA-A1 and EDA-A2, result in X-linked hypohidrotic ectodermal dysplasia (XLHED), which is characterized by defects in two or more ectodermal appendage types. EDA-A1 and EDA-A2 signal through the receptors EDAR and XEDAR, respectively. While the contributions of the EDA-A1/EDAR signaling pathway to ectodermal appendage phenotypes have been extensively characterized, the significance of the EDA-A2/XEDAR branch of the pathway has remained obscure. Herein, we report the phenotypic consequences of disrupting the EDA-A2/XEDAR pathway on mammary gland differentiation and growth. Using a mouse *Xedar* knock-out model, we show that *Xedar* has a specific and temporally restricted role in promoting post-pubertal growth and branching of the mammary epithelium that can be influenced by genetic background. Our findings are the first to implicate *Xedar* in ectodermal appendage development and suggest that the EDA-A2/XEDAR signaling axis contributes to the etiology of *EDA*-dependent mammary phenotypes.

## INTRODUCTION

The Ectodysplasin signaling pathway has long been recognized for its pivotal role in the development, pattering and differentiation of mammalian ectodermal appendages including hair follicles, eccrine sweat glands, teeth, and mammary glands (Biggs and Mikkola 2014; Cui et al. 2014; Cui et al. 2009; Headon et al. 2001; Headon and Overbeek 1999; Lindfors et al. 2013; Tucker et al. 2000; Voutilainen et al. 2015; Voutilainen et al. 2012; Wahlbuhl et al. 2018; Wahlbuhl-Becker et al. 2017). Alternative splicing of transcripts encoded by the anhidrotic ectodermal dysplasia gene (*Eda*) produces two main protein isoforms, EDA-A1 and EDA-A2, which belong to the tumor necrosis factor ligand superfamily (Yan et al. 2000). An insertion of two amino acids differentiates the EDA-A1 isoform from EDA-A2, and is necessary and sufficient to confer exclusive binding to the receptors EDAR and XEDAR, respectively (Yan et al. 2000). EDAR and XEDAR are type III transmembrane receptors whose respective binding to EDA-A1 and EDA-A2 oligomers has been shown to activate downstream NFκB signaling, and in some contexts JNK signaling (Kumar et al. 2001; Sinha et al. 2002; Yan et al. 2000).

In humans, loss of function mutations in *Eda* underlie the majority of cases of the most common form of ectodermal dysplasia, X-linked hypohidrotic ectodermal dysplasia (XLHED #MIM 305100) (Cluzeau et al. 2011). Affected individuals present with clinical features in two or more ectodermal appendages including reduced numbers or total loss of eccrine glands, sparse hair, missing teeth, and mammary phenotypes including impaired breast and nipple development, and lactation difficulties in females (Cluzeau et al. 2011; Wahlbuhl-Becker et al. 2017). The role of *Eda* in ectodermal appendage development is evolutionarily conserved. In *Tabbv* (*Ta*) mice, a base pair deletion in E*da* induces a frameshift resulting in non-functional EDA isoforms, causing highly homologous ectodermal appendage defects to those of human XLHED patients (Biggs and Mikkola 2014; Cui and Schlessinger 2006; Mikkola and Thesleff 2003; Sofaer and MacLean 1970; Srivastava et al. 1997; Wahlbuhl et al. 2018). XLHED-associated ectodermal appendage defects, particularly in hair follicles and eccrine glands, are thought to result from disruption of the EDA-A1/EDAR signaling axis, since loss of function mutations in *Edar* largely phenocopy the defects observed in *Eda^Ta^* mutants and exogenous treatment with recombinant EDA-A1 protein rescues or improves many of the XLHED hair and eccrine phenotypes in mice, humans, and dogs (Casal et al. 2007; Gaide and Schneider 2003; Margolis et al. 2019; Mustonen et al. 2004; Mustonen et al. 2003; Schneider et al. 2018; Srivastava et al. 2001). In contrast, the extent to which the EDA-A2/XEDAR signaling axis contributes to Eda-dependent ectodermal appendage phenotypes is unclear.

In humans, XLHED-inducing mutations generally result in dysfunction or loss of both EDA-A1 and EDA-A2 isoforms, as does the widely studied *Eda^Ta^* mouse mutation (Cluzeau et al. 2011; Wohlfart et al. 2016). Accordingly, deciphering the individual roles of the two ligand isoforms in ectodermal appendage biology relies on characterization of the effects of the receptors that mediate signaling by EDA-A1 and EDA-A2, respectively, namely EDAR and XEDAR. Unlike the dramatic ectodermal appendage phenotypes resulting from *Edar* loss-of function mutations, however, *Xedar* knock-out (*Xedar^KO^*) mice were reported to have normal hair, eccrine gland, and tooth development (Newton et al. 2004). Moreover, ectopic expression of an EDA-A2 transgene or recombinant Fc-EDA-A2 did not rescue hair, eccrine or tooth phenotypes in *Tabby* mice, underscoring the importance of the EDA-A1/EDAR pathway in the development of this subset of ectodermal appendages (Casal et al. 2007; Gaide and Schneider 2003; Margolis et al. 2019).

Intriguingly, analyses of the mammary glands of *Xedar^KO^* mice have not been reported. The mammary gland and its supporting structures are of clinical importance in humans since the majority of female XLHED carriers report lactation difficulties (Clarke et al. 1987). This is notable since treatment with or ectopic expression of EDA-A1 improved but did not fully rescue the mammary phenotypes of mouse *Tabby* mutants, including the reduction of mammary gland branching and lactation deficits (Mustonen et al. 2003; Srivastava et al. 2001; Wahlbuhl et al. 2018). In light of these observations and motivated by the clinical need to fully understand the etiology of human XLHED mammary phenotypes, we investigated the phenotypic consequences of disrupting the EDA-A2/XEDAR signaling axis on mammary gland morphogenesis using a constitutive *Xedar^KO^* mouse model.

## RESULTS AND DISCUSSION

### The loss of *Xedar* disrupts *Eda*-dependent epithelial mammary gland phenotypes

To establish a baseline spectrum of *Eda*-sensitive mammary phenotypes and to quantify the magnitude of effects of *Eda* loss on these traits, we evaluated two primary mammary traits previously reported to be attenuated in *Tabby* mutant females, namely, the size of the mammary gland (measured as the area invaded by the mammary epithelium into the mammary fat pad stroma), and the extent of branching of the mammary ductal tree (Chang et al. 2009; Voutilainen et al. 2012; Wahlbuhl et al. 2018), (Figure 1a-d). Analyses of the left and right, 4^th^ inguinal mammary glands of six-week-old, virgin female mice confirmed significant decreases in both gland area and branching in Eda^*Ta*^ homozygotes as compared to heterozygous and wildtype females on a C57BL/6J genetic background (Figure 1a-d and Table S1; χ^2^ =9.26(2), *P*<0.05; χ^2^ =17.9(2), *P*<0.05, respectively). Because mammary branching increases with body size in C57BL/6 sub-strains (Figure S1) and Tabby mice are smaller than their littermates (Table S1; χ^2^ =10.5(2), *P*<0.05), we confirmed that the disruption of branching that occurs with the loss of EDA signaling persists when branching is scaled by body size (Table S1; branches per gram; χ^2^ =17.6(2), *P*<0.05). The effect of *Eda* on mammary morphogenesis appears to be restricted to the epithelium as we did not find an effect of *Tabby* on the area of the mammary stroma, or fat pad (Tables S1, χ^2^ =2.31(2), *P*=0.315).

**Figure 1.**
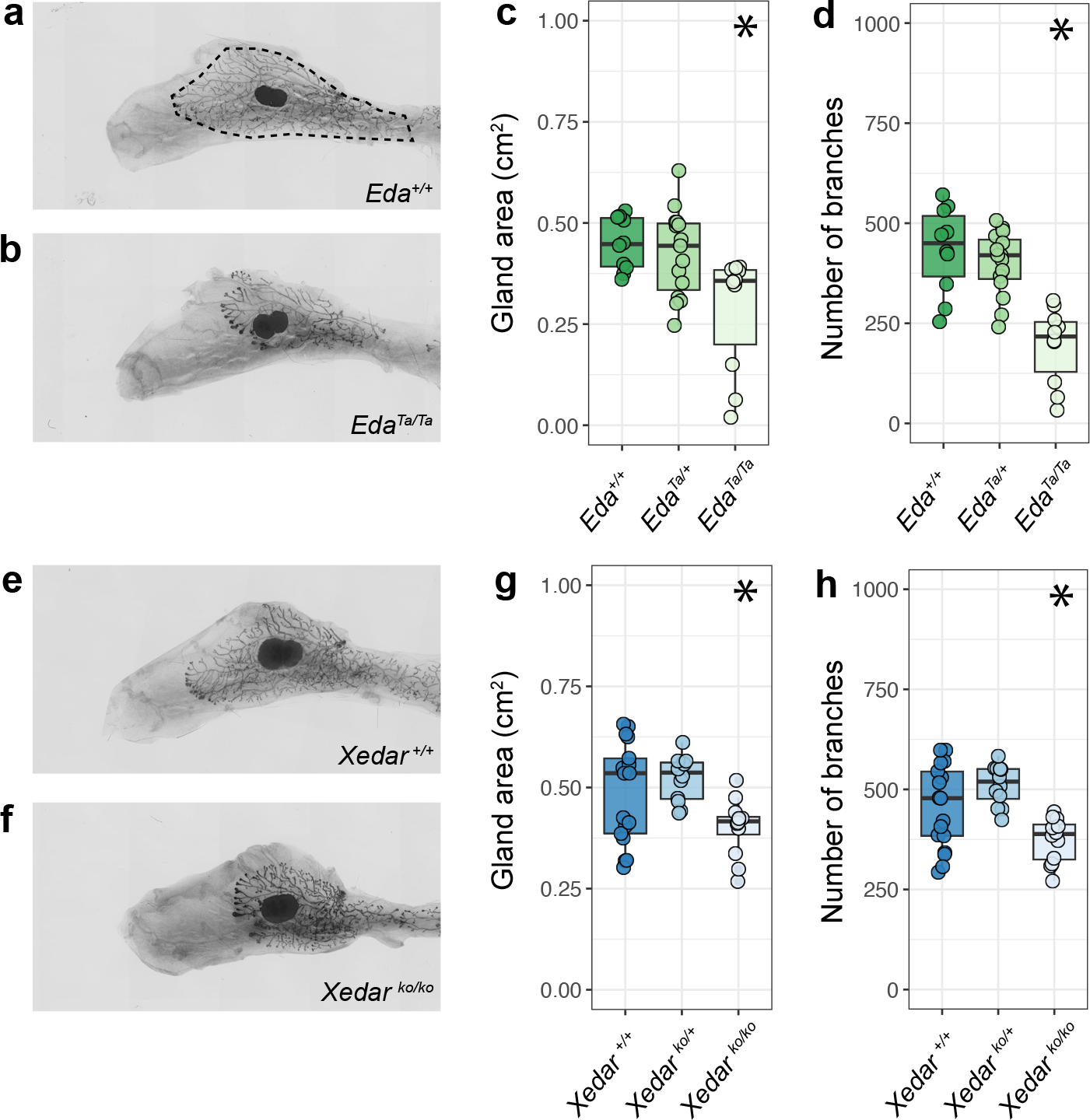
*Xedar* is required for development of *Eda*-dependent mammary epithelial traits. a,b. Representative images of the 4^th^ inguinal mammary gland from six-week-old, virgin female mice of designated *Eda* genotypes on C57BL/6J genetic background (+: wildtype allele; Ta: Tabby mutant allele). Dotted line indicates gland area. c. Area of the mammary epithelial tree across *Eda* genotypes. d. Epithelial branch count across *Eda* genotypes. e,f. Representative images of the 4^th^ inguinal mammary glands from six-week-old virgin mice of designated *Xedar* genotypes on C57BL/6N genetic background (KO: Knock Out allele). g. Area of the mammary epithelial tree across *Xedar* genotypes. h. Epithelial branch count across *Xedar* genotypes. Boxplots show median and quartile distributions for genotype categories. Dots represent phenotype values for individual mice analyzed in these experiments. Asterisks indicate *P*<0.05 by Kruskal Wallis testing.

Disruption of *Xedar* in C57BL/6N female mice affected both *Eda*-sensitive mammary gland traits. *Xedar^KO^* female mice exhibited reduced epithelial gland area and branching when compared to wildtype and hemizygous *Xedar^KO^* females (Figure 1e-h, Table S1: χ^2^ =9.41(2), *P*<0.05; χ^2^ =13.2(2), *P*<0.05, respectively). As with the loss of *Eda*, the effect of *Xedar* disruption was still observed when branching was scaled to body weight (Table S1: χ^2^ =18.5(2), *P*<0.05) and was restricted to the epithelium with no effect on fat pad area (Table S1: χ^2^ =0.19(2), *P*=0.906).

Since the C57BL/6J and C57BL/6N substrains show comparable baseline branching and gland size phenotypes (Figure 1a-h), we can qualitatively compare the effects of disrupting *Eda* and *Xedar* receptor on each of these mammary characteristics. Disrupting each gene results in a reduction of epithelial growth and branching. Branching is more severely attenuated with the loss of *Eda* than with the loss of *Xedar*, suggesting that *Eda* is able to support some branching in the absence of *Xedar*. Nevertheless, our data implicate *Xedar* in the differentiation of the adult mammary tree and provide the first direct evidence that *Xedar* affects *Eda*-dependent ectodermal appendage phenotypes.

### The effect of *Xedar* on mammary epithelial traits is dependent on genetic background

The patterns of growth and branching of the mammary epithelium vary among mouse strains (Gardner and Strong 1935; Naylor and Ormandy 2002). Nonetheless, the effect of *Eda* disruption via the *Tabby* allele has been reported across several mouse strains (Chang et al. 2009; Voutilainen et al. 2012). Indeed, we observe that disruption of *Eda* on an FVB/N strain background has consistent effects with those we observe in C57BL/6J mice (Table S1). To determine if *Xedar* loss showed similar phenotypic penetrance across genetic backgrounds, we examined the necessity of *Xedar* for normal mammary development in a second and genetically diverged laboratory mouse strain by backcrossing our *Xedar^KO^* allele onto FVB/N for at least four generations (N4), and compared the results to our C57BL/6N study (Lilue et al. 2018).

In six-week-old virgin N4 FVB/N female mice, mammary glands are larger and more branched than those of C57BL/6N mice even when accounting for the larger body size observed in the FVB/N strain (Figure S2, Table S1). Mammary gland area was significantly affected by *Xedar* genotype on an N4FVB/N background, with homozygous *Xedar^KO^* females showing a reduced gland area compared to wildtype or hemizygous carriers (Figure 2a-c and Table S1; χ^2^ =10.0(2), *P*<0.01). These data demonstrate that *Xedar* acts to promote the growth of the mammary epithelium on this genetic background much as is does in the C57BL/6N strain. In contrast, disruption of Xedar did not affect the number of mammary branches in FVB/N mice (Figure 2a, b, d and Table S1: gland area, χ^2^ =0.85(2), *P*=0.651).

**Figure 2.**
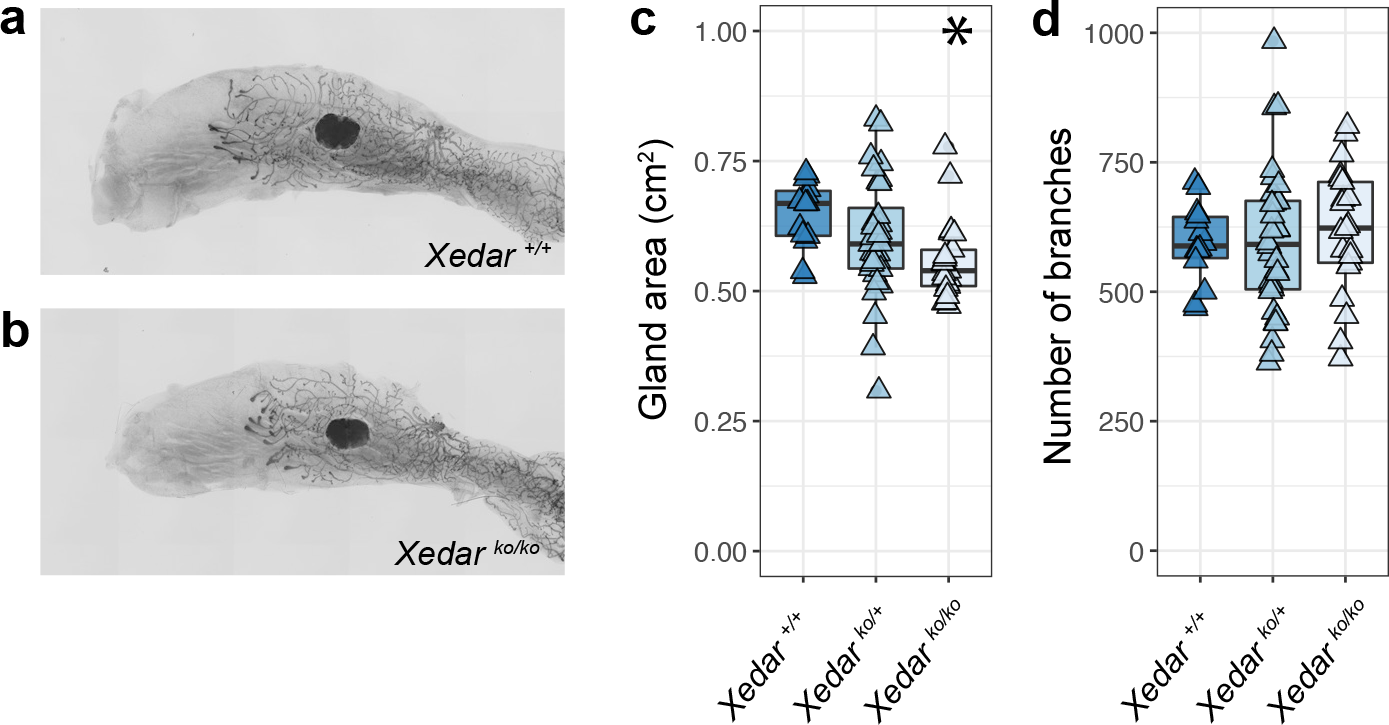
*Xedar* is necessary for post-pubertal mammary epithelial growth but not branching on an FVB/N genetic background. a,b. Representative images of the 4^th^ inguinal mammary gland of six-week-old, virgin, N4FVB/N female mice of the designated *Xedar* genotypes. c. Area of the mammary epithelial tree across *Xedar* genotypes (+: wildtype allele; KO: Knock Out allele). d. Number of epithelial branches within the mammary gland across *Xedar* genotypes. Boxplots show median and quartile distributions in each genotype category. Triangles represent phenotype values for individual mice analyzed in these experiments. Asterisks indicate *P*<0.05 by Kruskal Wallis tests.

*Xedar*’s ability to promote epithelial growth in two different mouse strains, but to promote branching in a strain-specific manner, suggests that genetic modifiers may influence the extent to which EDA-A2/XEDAR signaling contributes to *Eda*-dependent mammary phenotypes. These data demonstrate that underlying genetic context may have a profound influence on the phenotypic implications of *Xedar* variants, particularly in genetically diverse species such as humans.

### *Xedar* loss does not potentiate *Edar*-dependent mammary defects

In light of our discovery that *Xedar* impacts mammary phenotypes known to be sensitive to the EDA-A1/EDAR signaling axis, we investigated whether *Xedar* and *Edar* may independently or redundantly mediate the effects of *Eda* on mammary epithelial growth and branching. Consistent with previous reports, gland area and branch number were significantly reduced in the mammary glands of six-week-old virgin female mice homozygous for the *“Downless”(dl) Edar* loss-of-function mutation (Figure 3a-d; Table S1; χ^2^ =14.2(2), *P*<0.05; χ2 =18.5(2), *P*<0.05, for gland area and branching, respectively). Notably, *dl* heterozygotes are intermediate in branch number between wildtype controls and homozygotes, highlighting the sensitivity of this phenotype to EDA-A1/EDAR signaling (Figure 3d). This finding is consistent with the growing evidence that *Edar* is haploinsufficient for a subset of ectodermal appendage phenotypes (Kamberov et al. 2013).

**Figure 3.**
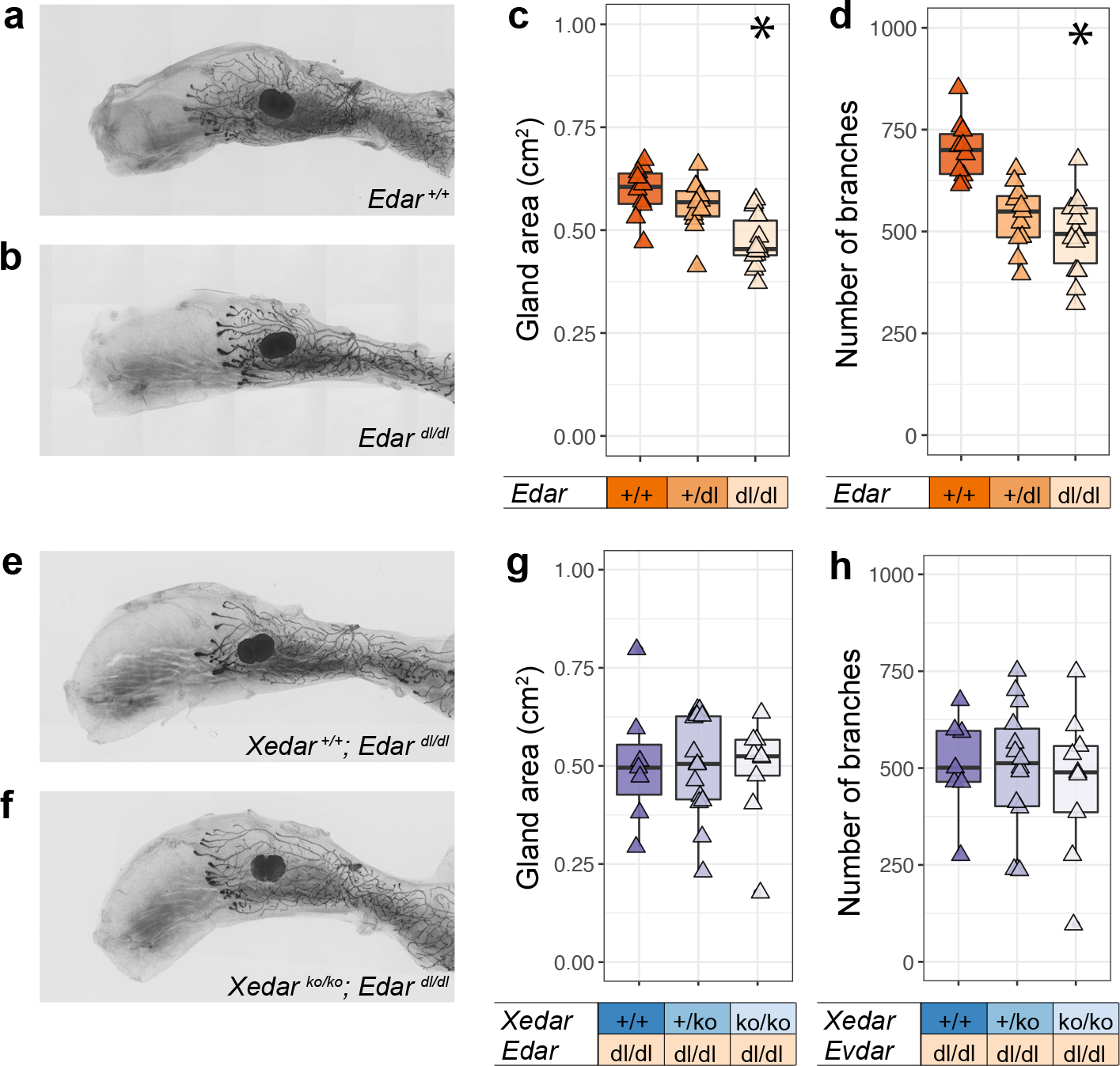
*Edar* is epistatic to *Xedar* in the regulation of post-pubertal mammary epithelium. a,b. Representative images of the 4^th^ inguinal mammary glands of six-week-old, virgin female mice of the designated *Edar* genotypes (+: wildtype allele; dl: *downless Edar* loss of function allele). c. Area of the mammary epithelial tree across *Edar* genotypes. d. Number of epithelial branches of the mammary gland across *Edar* genotypes. e,f. Representative images of the 4^th^ inguinal mammary glands of six-week-old, virgin female mice with compound disruptions in Edar and Xedar (KO: Knock Out allele). g. Area of the mammary epithelial tree across the designated *Edar* and *Xedar* compound genotypes. h. Number of epithelial branches of the mammary gland across the designated *Edar* and *Xedar* compound genotypes. Triangles represent phenotype values for individual mice analyzed in these experiments. Asterisks indicate *P*<0.05 by Kruskal Wallis tests.

By intercrossing *dl* mice with *Xedar^KO^* mice, we generated females that were homozygous for the *dl Edar* mutation and either wildtype, hemizygous or homozygous for the *Xedar^KO^* allele. Analysis of mammary gland area and branch number in these mice failed to reveal any effect of the loss of *Xedar* beyond the loss of *Edar* alone (Figure 3 e-h; Table S1; χ^2^ =0.22(2), *P*=0.892; χ^2^ =0.22(2), *P*=0.895).

The failure of *Xedar* disruption to potentiate *Edar-*dependent mammary phenotypes suggests that while *Xedar* and *Edar* can function independently, *Edar* is epistatic to *Xedar* in the regulation of mammary epithelial differentiation and growth. This may reflect a difference in the timing during which the two receptors regulate mammary morphogenesis or a difference in the downstream signaling mechanism engaged by each branch of the *Eda* pathway in this context. Previous studies have implicated NFκB as the major mediator of EDA-A1/EDAR signaling in mammary gland development, raising the possibility that in the context of the mammary gland, XEDAR functions through one of the alternative signaling mediators, such as JNK signaling (Kumar et al. 2001; Lindfors et al. 2013; Sinha et al. 2002; Voutilainen et al. 2015). Thus, our results suggest that multiple mechanisms may converge to regulate the mammary epithelial tree downstream of *Eda*.

### *Xedar* effects on mammary gland differentiation are temporally restricted

The development and differentiation of the mammary epithelium occurs in distinct stages beginning in mid-embryogenesis and continuing into adulthood (McNally and Martin 2011; Myllymäki and Mikkola 2019; Watson and Khaled 2008). Consistent with previous reports, we found that branching and size of the epithelial mammary tree is significantly reduced not only during the post-pubertal period prior to first estrus but also in pre-pubertal homozygous *Ta* mice at three weeks of age (Figure 4a,b: Gland area: χ^2^=10.2(2), *P*<0.05; branches: χ^2^=7.66(2), *P* <0.05) (Chang et al. 2009; Lindfors et al. 2013; Voutilainen et al. 2015; Voutilainen et al. 2012). In contrast, we did not observe significant differences in mammary gland area or epithelial branch number in three week old female mice carrying zero, one or two copies of the *Xedar^KO^* allele (Figure 4c,d: Gland area: χ^2^=2.15(2), *P* =0.341; branches: χ^2^=1.58(2), *P* =0.452). These data contrast with the effects of *Edar* disruption, which alters gland development at multiple stages including embryonically and during the pre-pubertal period (Lindfors et al. 2013; Voutilainen et al. 2015; Voutilainen et al. 2012). Instead, *Xedar*’s effects are temporally restricted to the post-pubertal stages of mammary gland development when hormonal cues provide the major directives for gland maturation. This is intriguing because we find that *Xedar* expression is detected both in the mammary epithelium as well as in the underlying mesenchyme during embryonic stages (Figure 4 e-h).

**Figure 4.**
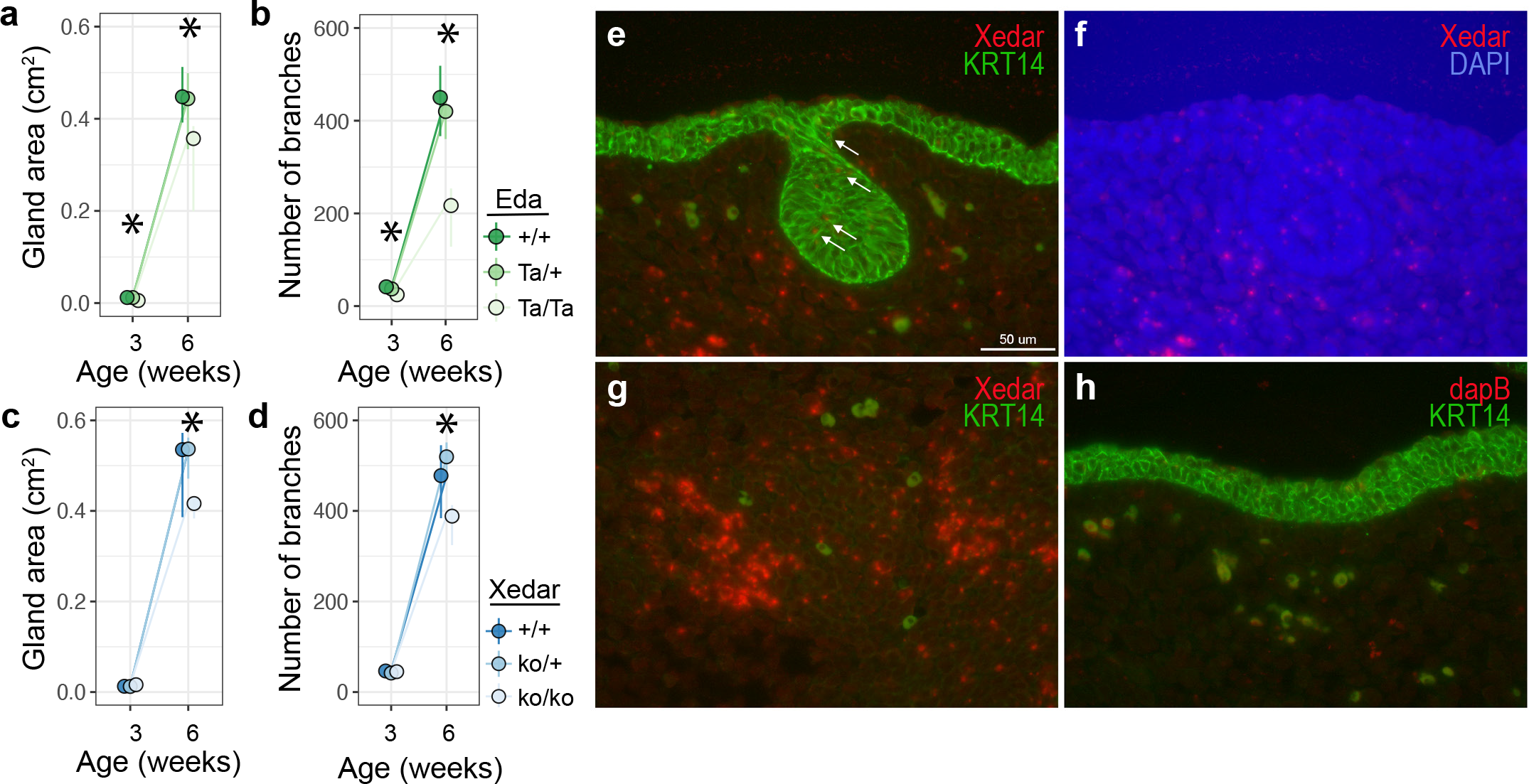
Effects of *Xedar* on mammary traits are restricted to post-pubertal period. a-d. Developmental assessment of 4^th^ inguinal mammary gland characteristics in three- and six-week-old virgin female mice. Area of the mammary epithelial tree (a) and branch count (b) differ across *Eda* genotypes (+: wildtype allele; Ta: Tabby mutant allele). Area of the mammary epithelial tree (c) and branch count (d) differ across *Xedar* genotypes at six but not three weeks (ko: knock out allele). Dots and whiskers show median and quartiles. Asterisks indicate *P*<0.05 by Kruskal Wallis tests. e,f,g. *In situ* studies showing expression of Xedar mRNA (red), Keratin 14 protein (KRT14, green), and 4′,6-diamidino-2-phenylindole (DAPI) nuclear stain (blue) in sections of mouse embryonic day 13.5 mammary bud (e,f) and muscle (g). White arrows: positive Xedar signal. h. *in situ* hybridization with negative control probe against dapB (red) in E13.5 skin section showing KRT14 positive basal keratinocytes.

The temporal restriction of the *Xedar^KO^* mammary phenotypes suggests a model in which *Xedar* and *Edar* differentially contribute to the regulation of mammary epithelial development and differentiation downstream of *Eda* at multiple stages of mammogenesis. In so doing, our findings point to a complex underlying basis for the effects of *Eda* on mammary glands. This may help to explain the incomplete rescue of mammary phenotypes in *Ta* mice by EDA-A1 alone and supports a contribution from the EDA-A2/XEDAR signaling axis in the etiology of mammary defects in human XLHED carriers. In a broader context, our study provides the first evidence implicating the EDA-A2/XEDAR signaling axis in the regulation of ectodermal appendage phenotypes. Unlike other characterized components of the *Eda* pathway, the effects of *Xedar* appear to be restricted to the mammary gland (Newton et al. 2004). Our finding that *Xedar*’s effects in the mammary gland can be sensitive to genetic background is noteworthy given that a derived *XEDAR* coding variant (XEDAR R57K, rs1385699) is highly differentiated among modern humans and was computationally identified as a potential target of positive natural selection in East Asia (Sabeti et al. 2007). Intriguingly, we have previously reported that a strongly selected coding variant of *EDAR* (EDARV370A, rs3827760) prevalent in present-day East Asian populations has pleiotropic effects on a subset of *EDA*-dependent ectodermal appendage traits, including on mammary gland branching (Kamberov et al. 2013). Thus, our findings raise the possibility that an additional *EDA* pathway effector, *XEDAR*, may also contribute to evolutionarily significant differences in mammary epithelial traits among modern humans.

## MATERIALS AND METHODS

### Experimental animals

Mice were housed in groups (up to 5 animals per cage) on a 12-hour light-dark cycle with continuous access to food and water. Pups were weaned at 3 weeks and raised thereafter in single sex groups. All experimental procedures were conducted in accordance with regulations and approvals by the Harvard Medical School and the Perelman School of Medicine Institutional Animal Care and Use Committees.

### Mouse lines

*Xedar* deficient mice (*Xedar* knock-out (*Xedar^KO^*)) mice have been previously described (Newton et al. 2004) and were obtained under material transfer agreement # OM-212731 from Genentech. *Xedar^KO^* mice harbor a targeted disruption in exon 4 of the *Xedar* locus leading to deletion of the XEDAR transmembrane domain and a non-functional protein (Newton et al. 2004). *Xedar^KO^* mice were obtained on a C57BL/6N genetic background and were maintained on C57BL/6N by further backcross to C57BL/6NTac (Taconic) mice. In addition, *Xedar^KO^* mice were separately backcrossed on to FVB/NCrl (Charles River) for at least four generations for analyses pertaining to effects of genetic background on mammary phenotypes. *Tabby* (*Ta*) mice (Jackson Labs, *A^w-J^*-*Eda^Ta-6J^*/J) harbor a loss of function mutation in *Ectodysplasin* (*Eda*) and were maintained on a C57BL/6J background (Jackson Labs C57BL/6J) and backcrossed onto FVB/NCrl (Charles River) for 4 generations to examine strain effects. *Downless (dl) Edar* loss of function mutant mice were obtained from Jackson labs (B6C3Fe *a*/*a*-*Edar^dl-J^*/J) and were backcrossed onto FVB/NCrl for at least eight generations prior to analyses.

To examine if *Xedar* deficiency could potentiate mammary phenotypes in the context of diminished EDAR signaling, we created mice with compound homozygous deficiency in *Xedar* and *Edar* on an FVB/N background. These lines were separately backcrossed to FVB/NCrl (Charles River) for four and eight generations, respectively, before the intercross. Experimental mice were either agouti or albino. All mice analyzed in the compound test crosses were *Edar*^*dl-J*/*dl-J*^ and showed symptoms of ectodermal dysplasia, including thin coat and a hairless, kinked tail.

### Genotyping

*Xedar^KO^* mice were genotyped using the following primers: Xedar-1 5’-tcgcaggactatgattgctaggc; Xedar-2 5’-gccatctgcatcaggtttcctatc; Xedar-3 5’-aggaaggcccattatcatgcagtc; Xedar-4 5’- ccagaggccacttgtgtagcg. The resulting PCR products are distinguishable by size using gel electrophoresis (wildtype band: 616 base pairs, mutant band: 302 base pairs).

*Dl* mice carry a loss of function mutation in ectodysplasin receptor that results in G/A substitution (5’gtgaaaacatggcgccaccttgcc G^(*wt*)^/A^(*dl-J*)^ agagctttggactgaag3’) in the *Edar* locus and an E379K amino acid change(Headon and Overbeek 1999). *dl* mutation was genotyped by sequencing of PCR product using the following primers (dl(J)-F 5’- gtctcagccccaccgagttg; dl(J)-R 5’- gtggggaggcaggtggtaca) to amplify genomic DNA from mouse tail biopsies. Tabby homozygote (*Eda*^*Ta-6J*/*Ta-6J*^), hemizygote (*Eda*^wt/*Ta-6J*^), and wildtype mice can be readily distinguished by eye when bred on pigmented strain. Homozygotes exhibit ectodermal dysplasia hair phenotypes and heterozygotes have a striped “tabby” coat. To allow our *Eda^Ta-6J^* FVB heterozygotes be phenotyped by eye (which is not possible in an albino), we maintained the line with an *A^w-J^* agouti allele.

### Tissue preparation

The 4^th^ and 5^th^ inguinal mammary glands and associated fat pad were dissected from 3 week and 6 week-old virgin female mice. Whole mount mammary preparations were made as follows: glands were fixed flat in Carnoy’s fixative (6 parts ethanol, 3 parts chloroform, 1 part glacial acetic acid) for 2 hours at room temperature and stored in 70% ethanol. Following rehydration, glands were stained overnight with Carmine alum solution (1g carmine Sigma C1022 with 2.5g aluminum potassium sulfate Sigma A7167 to 500ml with distilled water, boiled and filtered). Stained glands were dehydrated, cleared in xylenes, flat-mounted on glass slides, and imaged in brightfield with an Olympus VS120 slide scanner microscope.

### Analysis of mammary phenotypes

Mammary phenotypes were assessed using digital images analyzed in FIJI (NIH/ImageJ) with the Bioformats importer. Automatic branch counting was tested but was not as accurate as manual counting. Therefore ductal termini (branch tips) were counted manually using the FIJI Cell Counter plugin. Images were blind analyzed at least two times to ensure accuracy and reproducibility of measurements. Fat pad area was measured from the main lactiferous duct to the dorsolateral border. Gland length was measured from the distal-most ductal termini at the dorsal and ventral edges of the gland, capturing the maximum bidirectional growth of ductal tissue area across the fat pad. Gland area was measured by capturing the area invaded by the mammary epithelium from branch tip to tip across the extent of the mammary fat pad (see Figure 1). Left and right (4^th^ and 5^th^) gland counts and measurements were averaged for each individual in all experiments except the *Xedar-Edar* compound cross and the developmental series for which only left glands were used.

### Statistics

Statistical analyses were performed in R Statistical Software (v4.1.2; R Core Team 2021).). Mammary characteristics were compared using Kruskal-Wallis non-parametic tests, as normalcy requirements of parametric analysis were not met for all distributions in the dataset. Parametric and non-parametric analyses gave the same qualitative results in 94.7% of tests performed.

A minimum of ten females from each genotype class were used in all comparisons except the compound *Xedar-Edar* experiment for which only 7 *Xedar* wildtype and 9 *Xedar^KO/KO^; Edar^dl/dl^* compound mutant animals could be acquired. For our strain comparison, wildtype animals from all mouse lines were combined, resulting in 27 C57BL/6N and 34 FVB/N wildtype animals. This information is available in Table S1.

### *in situ* hybridization

CD1/NCrl (Charles River) embryos were harvested on embryonic day 13.5 (E13.5), fixed overnight with 4% paraformaldehyde (PFA) in 1× phosphate-buffered saline (PBS) and cryo-sectioned at 10 μm. RNAscope assay was performed as per manufacturer’s instructions for fixed frozen tissue and using RNAscope 2.5 Chromogenic assay reagent kit (Advanced Cell Diagnostics (ACD), United States). The *Xedar* (ACD: 531871), and the negative control dapB (ACD: 310043) probes were designed by ACD. Targeted regions for *Xedar* and dapB probes were 304 base pairs (bp) – 1253 bp and 414bp – 862bp in the transcript, respectively. Immunofluorescent staining to detect Keratin 14 was performed after RNAscope detection on the same tissue sections. The Keratin 14 antibody detects the basal keratinocyte layer of the ectoderm including the cells of the developing mammary gland which are also derivatives of this layer. Histological sections were prepared through the developing mammary bud and through muscle, in which Xedar was previously reported to be expressed. Briefly, samples were washed in PBS and blocked in PBS + 0.1% Tween (PBST) + 10% normal donkey serum before an overnight incubation in Cytokeratin 14 primary antibody (PRP155-P CK14, 1:10000, Covance). Samples were washed in PBST and incubated with Alexa Fluor^488^(1:250, Jackson ImmunoResearch) and 4′6-diamidino-2-phenylindole (1:5000, Sigma Aldrich). Images were acquired on a Leica DM5500 microscope equipped with a Leica DEC500 camera.

## DATA AVAILABILITY STATEMENT

All data analyzed in this study are reported in the manuscript. All mouse strains analyzed in this study have been previously published and are available from the vendors listed in the Material and Methods section of this manuscript.

## ACKNOWLEDGEMENTS

We thank Clifford J. Tabin (C.J.T.) for his support of this study. We also thank Joseph Patrice and Parimal Rana for excellent animal care. We thank Genentech for sharing the *XedarKO* mouse model (#OM-212731). We thank the Penn Biology and Diseases Resource-based Center Core A (P30-AR069589). The research reported in this publication was supported by a National Institute of Arthritis Musculoskeletal and Skin Diseases of the NIH Award R01AR077690 and a National Science Foundation BCS-1847598 award to Y.G.K.; and by a Eunice Kennedy Shriver National Institute of Child Health & Human Development of the NIH Award R01HD032443 to C.J.T.. Imaging was conducted at the Harvard Medical School NeuroImaging Facility with support from the Neural Imaging Center as part of an NINDS P30 Core Center Grant (P30-NS072030). Any opinions, findings and conclusions or recommendations expressed in this material are those of the authors and do not necessarily reflect the views of the National Science Foundation. The content is solely the responsibility of the authors and does not necessarily represent the official views of the National Institutes of Health.

## AUTHOR CONTRIBUTIONS STATEMENT

Conceptualization: AW, YGK; Data Curation: AW; Formal Analysis: AW, RT, DA, BW; Funding Acquisition: YGK; Investigation: AW, RT, DA, BK, BW; Project Administration: AW, YGK; Supervision: AW, YGK; Validation: AW, RT; Visualization: AW, DA, BK; Writing - Original Draft Preparation: AW; Writing - Review and Editing: AW, YGK

