## Supplemental Material for "Ectodysplasin signaling via Xedar is required for mammary gland morphogenesis"

### SUPPLEMENTARY MATERIALS_Wark et al 2022

**Table S1. Comparison of mammary characteristics of adult female mice by genotype.**

| Gene (Allele) | Strain | | N | Weight (g) | Gland area ($cm^2$) | Number of branches | Branches per gram (#/g) | Fat pad area ($cm^2$) |
|---|---|---|---|---|---|---|---|---|
| *Eda (Tabby)* | C57BL/6J | wt | 10 | 17.1 ± 1.5 | 0.45 ± 0.07 | 433.4 ± 108.1 | 25.1 ± 5.1 | 0.79 ± 0.11 |
| | | het | 15 | 16.8 ± 1.0 | 0.43 ± 0.11 | 400.4 ± 79.3 | 23.6 ± 4.0 | 0.73 ± 0.15 |
| | | hom | 10 | 15.0 ± 1.2 | 0.28 ± 0.15 | 194.0 ± 95.2 | 12.5 ± 5.8 | 0.70 ± 0.14 |
| | | $\chi^2$ | | 10.5(2) | 9.26(2) | 17.9(2) | 17.6(2) | 2.31(2) |
| | | *P* | | **0.005** | **0.010** | **1.3E-04** | **1.5E-04** | 0.315 |
| *Xedar (KO)* | C57BL/6N | wt | 17 | 16.8 ± 1.3 | 0.49 ± 0.12 | 456.4 ± 103.9 | 26.9 ± 4.3 | 0.75 ± 0.11 |
| | | het | 12 | 17.2 ± 1.1 | 0.52 ± 0.06 | 511.2 ± 50.2 | 29.6 ± 1.8 | 0.76 ± 0.08 |
| | | hom | 12 | 17.2 ± 1.5 | 0.40 ± 0.07 | 373.7 ± 55.6 | 21.7 ± 2.4 | 0.74 ± 0.12 |
| | | $\chi^2$ | | 0.55(2) | 9.41(2) | 13.2(2) | 18.5(2) | 0.19(2) |
| | | *P* | | 0.758 | **0.009** | **0.001** | **9.26E-05** | 0.906 |
| *Eda (Tabby)* | N5FVB/N | wt | 10 | 22.1 ± 2.9 | 0.59 ± 0.06 | 770.8 ± 133.3 | 34.9 ± 4.8 | 0.98 ± 0.13 |
| | | het | 12 | 20.3 ± 1.8 | 0.55 ± 0.08 | 592.8 ± 90.1 | 29.1 ± 3.8 | 0.98 ± 0.11 |
| | | hom | 10 | 18.9 ± 1.8 | 0.38 ± 0.13 | 434.0 ± 183.1 | 22.5 ± 8.5 | 0.85 ± 0.14 |
| | | $\chi^2$ | | 6.29(2) | 15.8(2) | 15.2(2) | 13.2(2) | 5.75(2) |
| | | *P* | | **0.043** | **3.67E-04** | **4.81E-04** | **0.001** | 0.056 |
| *Xedar (KO)* | N4FVB/N | wt | 14 | 21.1 ± 1.9 | 0.65 ± 0.06 | 593.2 ± 75.7 | 28.3 ± 4.6 | 1.09 ± 0.16 |
| | | het | 32 | 19.6 ± 1.7 | 0.60 ± 0.11 | 597.6 ± 143.1 | 30.5 ± 7.2 | 0.94 ± 0.16 |
| | | hom | 21 | 19.7 ± 1.4 | 0.56 ± 0.08 | 618.7 ± 123.1 | 31.4 ± 6.0 | 0.85 ± 0.12 |
| | | $\chi^2$ | | 5.60(2) | 10.0(2) | 0.85(2) | 3.00(2) | 16.6(2) |
| | | *P* | | 0.061 | **0.007** | 0.651 | 0.222 | **2.5E-04** |
| *Edar (Downless)* | N7FVB/N | wt | 10 | 22.3 ± 2.8 | 0.59 ± 0.06 | 699.2 ± 74.0 | 31.7 ± 4.4 | 0.88 ± 0.09 |
| | | het | 11 | 20.6 ± 1.4 | 0.56 ± 0.06 | 535.1 ± 79.8 | 25.9 ± 4.2 | 0.87 ± 0.07 |
| | | hom | 14 | 18.8 ± 1.0 | 0.47 ± 0.06 | 492.2 ± 95.6 | 26.1 ± 4.4 | 0.86 ± 0.13 |
| | | $\chi^2$ | | 19.8(2) | 14.2(2) | 18.5(2) | 8.62(2) | 0.14(2) |
| | | *P* | | **4.8E-05** | **0.001** | **9.40E-05** | **0.013** | 0.931 |
| *Xedar (KO)* | N4FVB/N; *Edar* $^{dl/dl}$ | wt | 7 | 19.2 ± 1.6 | 0.50 ± 0.16 | 510.2 ± 130.0 | 26.6 ± 6.8 | 0.88 ± 0.12 |
| | | het | 14 | 19.6 ± 1.2 | 0.50 ± 0.13 | 489.3 ± 176.7 | 24.7 ± 8.2 | 0.89 ± 0.08 |
| | | hom | 9 | 19.3 ± 1.4 | 0.48 ± 0.13 | 464.5 ± 191.8 | 23.7 ± 9.7 | 0.92 ± 0.11 |
| | | $\chi^2$ | | 0.21(2) | 0.22(2) | 0.22(2) | 0.47(2) | 0.63(2) |
| | | *P* | | 0.897 | 0.892 | 0.895 | 0.789 | 0.727 |

Means +/- standard deviations shown with Kruskal Wallis test statistics. Signficant effects are bold-faced. KO=knockout; dl=downless.

**Figure S1. Mammary branch number increases with body weight in substrains of C57BL/6.** Epithelial branch number is plotted against body weight for individual wildtype adult virgin females at 6 weeks of age from C57BL/B6N and C57BL/6J mouse substrains. Dots represent phenotype values for individual mice analyzed in these experiments.

**Figure S2. Bodyweight and mammary characteristics are differentiated between mouse C57BL/6 substrain and FVB/N genetic backgrounds.** a-d. Characteristics of six-week-old virgin wildtype female mice from two genetic background categories: C57BL/6 (including

C57BL/6N and C57BL/6J substrains) and FVB/N (including individuals that are four to eight generations backcrossed to FVB/N). Dots and triangles represent phenotype values for individual mice analyzed in these experiments, including bodyweight in grams (a), mammary epithelial tree area (b), number of epithelial branches (c), and number of branches scaled to body weight (d). Boxplots show median and quartile ranges for each characteristic. Asterisks indicate $P<0.05$ by Kruskal Wallis tests.

Figure S1

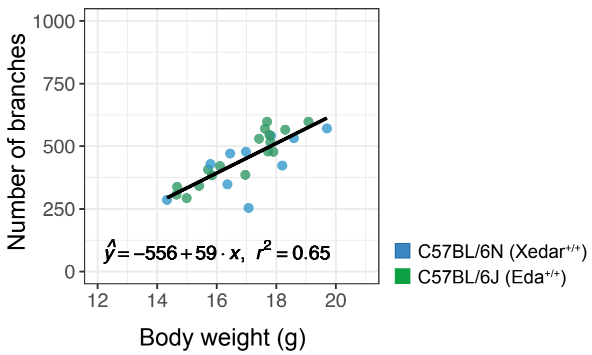

Number of branches
1000
750
500
250
0
12
14
16
18
20
Body weight (g)
$\hat{y} = -556 + 59 \cdot x,\ r^2 = 0.65$
C57BL/6N (Xedar+/+)
C57BL/6J (Eda+/+)

Figure S2

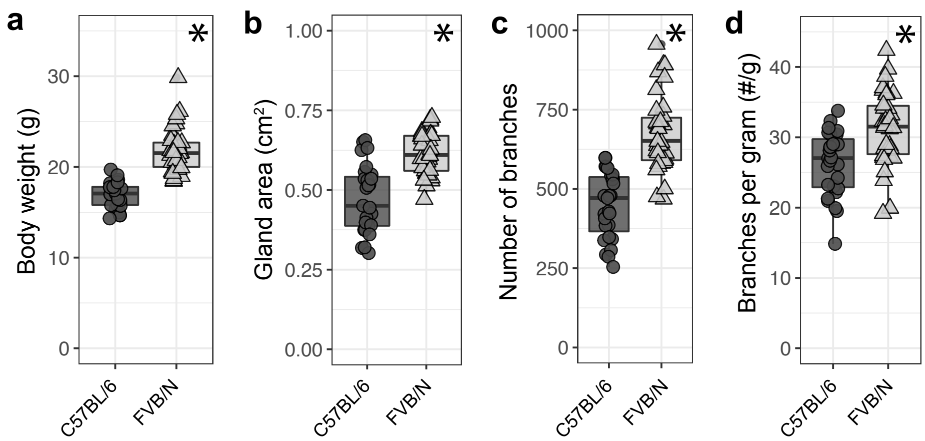
a
b
c
d
Body weight (g)
Gland area (cm²)
Number of branches
Branches per gram (#/g)
C57BL/6
FVB/N
*
